# Addressing the antibody germline bias and its effect on language models for improved antibody design

**DOI:** 10.1101/2024.02.02.578678

**Authors:** Tobias H. Olsen, Iain H. Moal, Charlotte M. Deane

## Abstract

The versatile binding properties of antibodies have made them an extremely important class of biotherapeutics. However, therapeutic antibody development is a complex, expensive and time-consuming task, with the final antibody needing to not only have strong and specific binding, but also be minimally impacted by any developability issues. The success of transformer-based language models in protein sequence space and the availability of vast amounts of antibody sequences, has led to the development of many antibody-specific language models to help guide antibody discovery and design. Antibody diversity primarily arises from V(D)J recombination, mutations within the CDRs, and/or from a small number of mutations away from the germline outside the CDRs. Consequently, a significant portion of the variable domain of all natural antibody sequences remains germline. This affects the pre-training of antibody-specific language models, where this facet of the sequence data introduces a prevailing bias towards germline residues. This poses a challenge, as mutations away from the germline are often vital for generating specific and potent binding to a target, meaning that language models need be able to suggest key mutations away from germline.

In this study, we explore the implications of the germline bias, examining its impact on both general-protein and antibody-specific language models. We develop and train a series of new antibody-specific language models optimised for predicting non-germline residues. We then compare our final model, AbLang-2, with current models and show how it suggests a diverse set of valid mutations with high cumulative probability. AbLang-2 is trained on both unpaired and paired data, and is freely available (https://github.com/oxpig/AbLang2.git).

## 1 Introduction

The potential of antibodies, also known as B-cell Receptors (BCR), to bind and neutralise any pathogen by either blocking their function or marking them for removal [1, 2, 3], has made them a valuable tool in many areas of medical and scientific research. Antibodies are used routinely in diagnostic assays [4] and are by far the most successful class of biotherapeutic, evidenced by the more than 170 antibody therapeutics that are in regulatory review or approved for clinical use to date [5, 6]. However, therapeutic antibody development is a complex task, traditionally taking years. To be a therapeutic, an antibody needs to not only bind strongly and exclusively to its target, but also be minimally impacted by undesirable properties, like aggregation [7], damaging post-translational modification sites [8], and immunogenicity [9, 10], collectively known as developability issues [11]. Antibodies therefore go through multiple rounds of optimization, leading to an expensive and time-consuming development process. As a consequence, there has been an increasing focus on developing computational methodologies to help this development [3].

Recently, transformer-based language models (LMs) have become indispensable for the prediction of many language tasks, like language translation, question answering and human-like text generation [12, 13, 14, 15]. Given the shared similarities between protein sequences and natural language, both comprised of basic units in the form of amino acids and words with an inherent syntax, a lot of effort has been put into developing and training protein-specific LMs (e.g. [16, 17, 18, 19, 20]) and for antibody discovery and design, antibody-specific LMs (e.g. [21, 22, 23, 24]). The key element for LMs is that they are initially pre-trained on vast amounts of unlabelled data in an unsupervised manner [12].

Pre-trained LMs can then be used to create information-rich representations of the input sequence or, as popular for BERT-like protein and antibody LMs, for directed evolution by suggesting mutations that lead to better binding and developability properties [25]. BERT-like models are inspired by the BERT architecture and trained using the masked language modeling (MLM) approach [12].

Most antibody-specific LMs are pre-trained on the antibody sequences in databases like the Observed Antibody Space (OAS) [26]. These sequences are obtained through high-throughput BCR repertoire sequencing (BCR-seq), which enables sequencing of millions of antibodies per sample [27]. However, because of limitations on the possible length that can be sequenced, only the variable domain of the heavy (VH) or light (VL) chain is usually sequenced, as the VH and VL contain most of the sequence diversity and the binding site [28]. The complete binding site spans both the VH and VL, but BCR-seq methods retaining the VH-VL pairing information currently have a much lower throughput [29]. Thus, most antibody-specific LMs have been trained purely on unpaired VH and VL sequences, with only a few exceptions [30].

Each VH and VL consists of a framework region (FWR) and three loops, called complementarity-determining regions (CDR)1-3. These three CDRs on each chain also constitute the majority of the binding site [2, 31]. Antibody diversity is primarily located in the CDRs, with the CDR3 being especially diverse because of the combination of different gene segments during V(D)J recombination. The VL is comprised of a variable (V) segment and a joining (J) segment, with its CDR3 spanning both, while the VH also contains a diversity (D) segment fully contained within the CDR3 [2, 31]. During infection, non-germline (NGL) mutations are introduced via somatic hypermutation (SHM), to develop strong target-specific binding [2, 31]. However, the majority of the sequence of these affinity-matured antibodies are still germline [31]. Moreover, BCR-seq is often performed on blood samples as these are less invasive to obtain compared to other samples [27]. Blood is characterised by a low proportion of affinity-matured antibody producing B-cells, such as memory B-cells and plasma B-cells. Thus, BCR-seq often yields antibodies mostly from naive B-cells, which have not yet undergone SHM [27]. The data used to train antibody-specific LMs is therefore likely to be heavily biased towards the germline.

LMs are known to reproduce and even amplify biases in their training data [32]. Protein LMs are similarly affected, having been shown to struggle with mutations far from the wildtype [33]. For natural language LMs, efforts to reduce biases have included pre-processing training data [32] or de-biasing with fine-tuning [34], while recalibration for each individual protein with respect to the background distribution of random mutation has been tried for protein LMs [33]. The germline bias in antibody sequences can also be viewed as an imbalance problem. When predicting randomly selected masked residues, it is rarely a NGL residue which needs to be predicted. The imbalance problem is well-known and many solutions have been proposed, like up or down-sampling [35] and focal loss [36]. Focal loss is a loss function that down-weights the loss of well predicted labels. As rare labels, such as NGL residues, are usually poorly predicted, it results in an increased focus on these labels during training.

While affinity-matured antibodies usually only contain a few NGL mutations outside the CDR3, their existence is often important for specific and high-affinity binding [31]. It is therefore necessary to understand if and how the germline bias affects antibody-specific LMs, especially their ability to suggest relevant NGL mutations. Correctly selecting relevant NGL residues might result in the design and optimization of better therapeutic antibodies than the current protein and antibody-specific LMs.

In this study, we first explore the germline bias in antibody sequences, both from BCR-seq data and a set of therapeutic antibodies. We then investigate how the bias affects BERT-like LMs’, like ESM-2 [18] and various antibody-specific LMs, ability to predict NGL residues. We then iteratively train and improve a new antibody-specific LM specifically for NGL prediction, and show how our final model, AbLang-2, is able to more accurately suggest a diverse set of valid mutations compared to previous models.

## 2 Methods

### 2.1 Dataset preparation

The training and test sets were derived from the Observed Antibody Space (OAS) [26]. Antibody sequences were downloaded from OAS in Nov. 2022, yielding 2,072M VHs, 357M VLs and 1.57M paired antibodies. The sequences were then filtered by removing duplicates, sequences missing conserved cysteines, and heavily fragmented (missing more than 16 residues from the N-terminus or 7 residues from the C-terminus) sequences. Unpaired sequences were additionally filtered to remove sequences only seen once. Finally, any amino acids other than the standard 20 were changed to X.

Redundancy was then reduced with clustering. Unpaired sequences were clustered first based on identical CDR3s and thereafter by 95% identity over the whole sequence using Linclust [37] with –cov-mode 1. This clusters fragments together with a longer representative sequence. Paired sequences had first their VH and VL clustered individually as done for unpaired sequences. The paired sequences were then clustered by having the same VH and VL cluster. The longest sequence or sequence pair from each cluster was kept.

The paired antibodies were then randomly split into a train and test set of 1.26M and 100k, respectively. The paired test set was clustered together with the reduced unpaired set by 95% identity over the whole sequence using Linclust [37] with –cov-mode 1. Any unpaired sequences clustered with a VH or VL from the paired test set, were removed. This resulted in training sets with 27.5M VHs, 11.1M VLs and 1.26M paired antibodies, and a test set of 100k paired antibodies.

The therapeutic sequences used in this study were sourced from Thera-SAbDab (as of Feb. 2023) [6]. Only VH-VL paired antibodies were selected, resulting in 735 therapeutic test cases.

### 2.2 Germline and non-germline residue estimation

For OAS derived sequences, germline and NGL residues were determined with IgBLAST [38]. IgBLASTn uses the nucleotide sequences to predict each antibody’s germlines, including non-templated regions within the CDR3, which was then used to label each residue. For the 735 therapeutic sequences, the germlines were predicted with ANARCI [39] from the protein sequence. The ANARCI predicted germline sequences were then used to label each residue as germline or NGL.

The non-templated regions in the VH and VL CDR3s, and the uncertainty in estimating the D germline within the VH CDR3, complicates the identification of germline residues within the CDR3s. For both the VH and VL CDR3s, we therefore only measured the NGL perplexity, which has the estimated germline residues, including the estimated D germline for the VH CDR3, filtered away.

For a clear separation of germline and NGL residues, we focus on mutations outside of the CDR3. While estimating the V and J gene using IgBLAST is relatively reliable, it depends on a database of known V and J genes. The potential lack of certain alleles in this database, could result in the misclassification of some germline residues as NGL. Nonetheless, with most obvious germline residues filtered out, this approach will result in a much more challenging dataset than random selection.

The standard, germline, and NGL residue test sets were generated from sequences in the test set (see Methods 2.1). The standard test set represents how perplexity is usually measured, and is a random selection of 20,000 residues. For the germline test set, we sampled 20,000 estimated germline residues from outside the CDR3. For the NGL test set, we used all 475,000 NGL residues outside the CDR3 found in the test set, and a random selection of 9,000 NGL residues within the CDR3.

### 2.3 Perplexity calculation

Perplexity is commonly used for measuring the uncertainty of LM predictions and for performance comparison. For BERT-like LMs, sequence perplexity can be derived by first computing the cross-entropy for each predicted masked residue individually. The sequence perplexity is then the exponential of the mean of these losses, and the perplexity of a test set is then the average sequence perplexity [40]. Instead of measuring perplexity based on every residue in a sequence, we measured perplexity based on a subset of residues across the test set. This subset can be only NGL or germline residues, or a combination as with the standard test set. For consistency, the same residues are used to assess the perplexity for each model.

### 2.4 Architecture and training

A series of models (see Table 1) were iteratively improved and trained using the training sets (see Methods 2.1). The models were implemented in PyTorch 2.0.1 [41] and trained using the PyTorch-Lightning framework [42]. The initial model (Ab-Unpaired) was based on the architecture of a 6-layered ESM-2 model with SwiGLU [43] as its activation function, and trained on single chains from the paired training set. The model was optimized with an Adam optimizer. For stabilizing and enhancing training, we used a linear warm-up for 1k steps, a peak learning rate of 0.0004, a cosine learning rate decay over 9k steps, and a weight decay of 0.01. An effective batch size of 8192 was used during the training, together with a layer normalization with an epsilon of 1e^*−*12^. Training occurred using the standard MLM training approach of randomly masking and predicting 15% of the input residues.

**Table 1:** Comparison of the architecture, training data, and training approach for the protein language model (LM) ESM-2 [18], the antibody-specific LMs AntiBERTy [22] and AbLang-1 [24], and our new selection of antibody-specific LMs. The architecture column shows the most similar architecture and the model’s size with the number of layers (L) and embedding size (ES). While the exact number of training steps for AntiBERTy is unknown, it was trained for 8 epochs [22]. AbLang-1 and the new antibody-specific LMs were trained on 8192 sequences (4096 for AbLang-1 Light) per batch, with each sequence comprising approximately 120 amino acids. Each batch thus contained about 1M tokens for unpaired sequences and 2M for paired antibody VH-VL sequences. CE, cross-entropy loss; FL, focal loss; MLM, masked language modeling

|  | Architecture | Training data | Paired | Loss function | Training objective | Training steps | Batch size |
| --- | --- | --- | --- | --- | --- | --- | --- |
| ESM-2 | ESM-2<br>33L + 1280ES | UR50/D<br>60M sequences | N | CE | MLM | 500K | 2M tokens |
| AntiBERTy | BERT<br>8L + 512ES | 558M VH/VL | N | CE | MLM | 8 epochs | N/A |
| AbLang-1 | RoBERTa<br>12L + 768ES | 187K VL<br>14.2M VH | N | CE | MLM | 2K<br>34K | 500K tokens<br>1M tokens |
| Ab-Unpaired | ESM-2<br>6L + 320ES | 1.26M VL<br>1.26M VH | N | CE | MLM | 10K | 1M tokens |
| Ab-Paired | ESM-2<br>6L + 320ES | 1.26M paired | Y | CE | MLM | 10K | 1-2M tokens |
| Ab-FL | ESM-2<br>6L + 320ES | 1.26M paired | Y | FL | MLM | 10K | 1-2M tokens |
| Ab-ModMask | ESM-2<br>6L + 320ES | 1.26M paired | Y | FL | Modified<br>MLM | 10K | 1-2M tokens |
| Ab-FT | ESM-2<br>6L + 320ES | 35.6M VH/VL<br>1.26M paired | Y | FL | Modified<br>MLM | 10K + 1K | 1-2M tokens |
| AbLang-2 | ESM-2<br>12L + 480ES | 35.6M VH/VL<br>1.26M paired | Y | FL | Modified<br>MLM | 200K + 10K | 1-2M tokens |

The model was then further improved over several iterations, with each new model being an expansion of the previous one. Ab-Paired: The input was modified to also handle paired antibodies, by separating true VH-VL pairs with a separator token. The model was then trained with unpaired VH and VL, and paired VH-VL chains, from the paired training set. Ab-FL: Instead of the conventional cross-entropy loss function, focal loss was used [36]. The purpose of this loss function is to better address the challenge of imbalanced or sparse datasets. Ab-ModMask: The standard MLM approach was modified to include two alternative masking methods; short 3-5 segment masking and singular large segment masking, both inspired by [44]. For each batch, a masking method (the two new masking methods and standard MLM) is then selected uniformly. The proportion of masked residues was also changed to a dynamic value, selected uniformly between 10% and 40%. Ab-FT: The model was initially pre-trained exclusively on the unpaired sequences (see Methods 2.1) for 10,000 steps. This was followed by fine-tuning on paired sequences for an additional 1,000 steps and a peak learning rate of 0.0001. AbLang-2: The architecture was scaled up to 12 layers and an embedding size of 480. The model was then pre-trained on unpaired sequences for 200,000 steps and subsequently fine-tuned for 10,000 steps on paired sequences.

## 3 Results

### 3.1 Germline bias in antibody sequence data

To investigate germline bias, we inspected the VH-VL sequences of all paired antibodies within OAS [26]. The majority of these antibodies originate from naive B-cells (42%) and unsorted B-cells (39%), with only 17% from memory B-cells. The last 1% of antibodies are derived from other cells like plasma B-cells (see Fig. 1a). The paired data in OAS is therefore predominantly derived from B-cells that have not undergone SHM.

**Figure 1:**
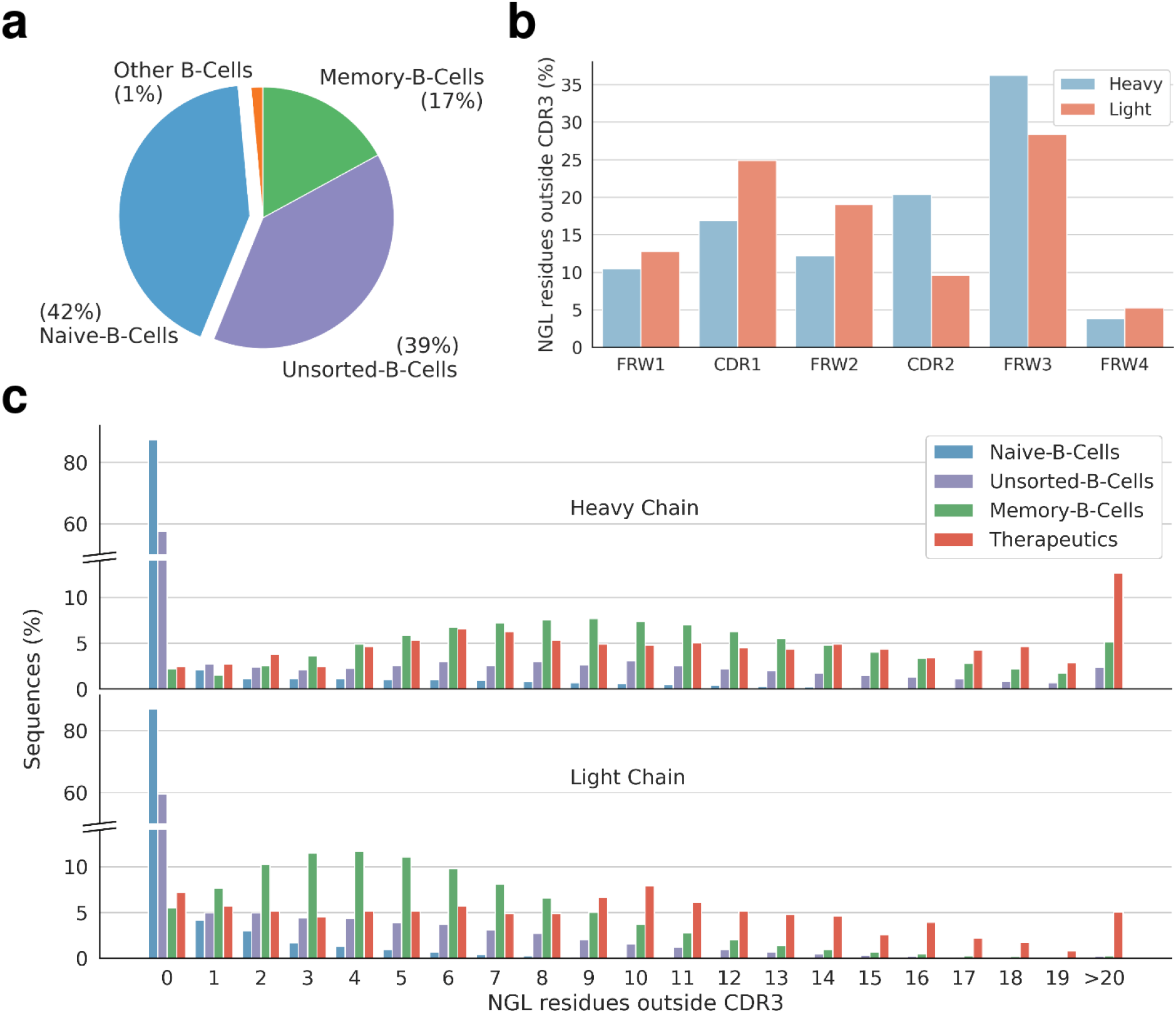
Overview of non-germline (NGL) residues outside of the CDR3 from paired antibody VH-VL sequences in OAS. **a**, Distribution of OAS derived antibody origins, showing naive B-cells as the predominant source (42%), followed by unsorted B-cells (39%) and memory B-cells (17%). **b**, Distribution of NGL residues across different regions. **c**, Distribution of NGL residues outside the CDR3 per sequence by source. Naive B-cell derived antibodies lack NGL residues, while memory B-cell derived antibodies display an average of *∼*10 and *∼*5.3 NGL residues in their VH and VL, respectively. Therapeutic antibodies exhibit averages of *∼*11.5 and *∼*8.8 NGL residues for the VH and VL. Supplementary Fig. 1S provides an extended view of the distribution across both chains, with memory B-cell and therapeutic antibodies averaging *∼*15.3 and *∼*20.3 NGL residues, respectively.

NGL residues outside the CDR3 were identified (see Methods 2.2) and their distribution across different regions (see Fig. 1b) and different cell sources (see Fig. 1c) compared. As expected, the majority of antibodies from naive B-cells lack NGL residues, while those from memory B-cells contain a large number of NGL residues, averaging *∼*10 and *∼*5.3 in the VH and VL, respectively. For comparison, a slightly higher count of NGL residues was observed for antibody therapeutics (see Methods 2.2), averaging *∼*11.5 and *∼*8.8 in the VH and VL, respectively. Supplementary Fig. S1 shows the distribution across both chains. Here, memory B-cell derived antibodies averaged *∼*15.3 NGL residues, while therapeutic antibodies showed an average of *∼*20.3 NGL residues.

### 3.2 Germline bias in pre-trained language models

The impact of the germline bias in antibody sequences on pre-trained LMs was investigated for a set of LMs, both general-protein (ESM-2 [18]) and antibody-specific (Sapiens [21], AntiBERTy [22] and AbLang-1 [24]) LMs. The ESM-2 models were trained on the UniRef50 [45] dataset, comprising approximately 60M protein sequences, of which a few hundreds are antibody sequences [18]. In this work we use the 650M-parameter ESM-2 model. The antibody-specific LMs are trained solely on unpaired VH and VL sequences from OAS. Sapiens was trained on 20M and 19M human VH and VL sequences, respectively [21]. AntiBERTy was trained on 558M VH and VL sequences from various species [22]. AbLang-1 was similarly trained on 14M and 187k VH and VL sequences from a mix of species [24].

The impact was first examined by investigating how often the germline is predicted for masked NGL residues. For this we used the previously derived NGL residues and predicted the masked residues with the four LMs. Fig. 2 shows how often the germline was predicted for both the VH and VL, with sequences grouped along the x-axis by their number of NGL residues. Sapiens, AntiBERTy, and AbLang-1, predicted the germline with frequencies of 87.6%, 86.7%, and 84.9%, respectively. ESM-2, despite its limited exposure to antibody sequences, still predicted the germline at a rate of 49.6%. While it remains unclear if models less germline biased are better at predicting NGL residues, it is clear that all the models tested preferentially suggest mutations towards the germline.

**Figure 2:**
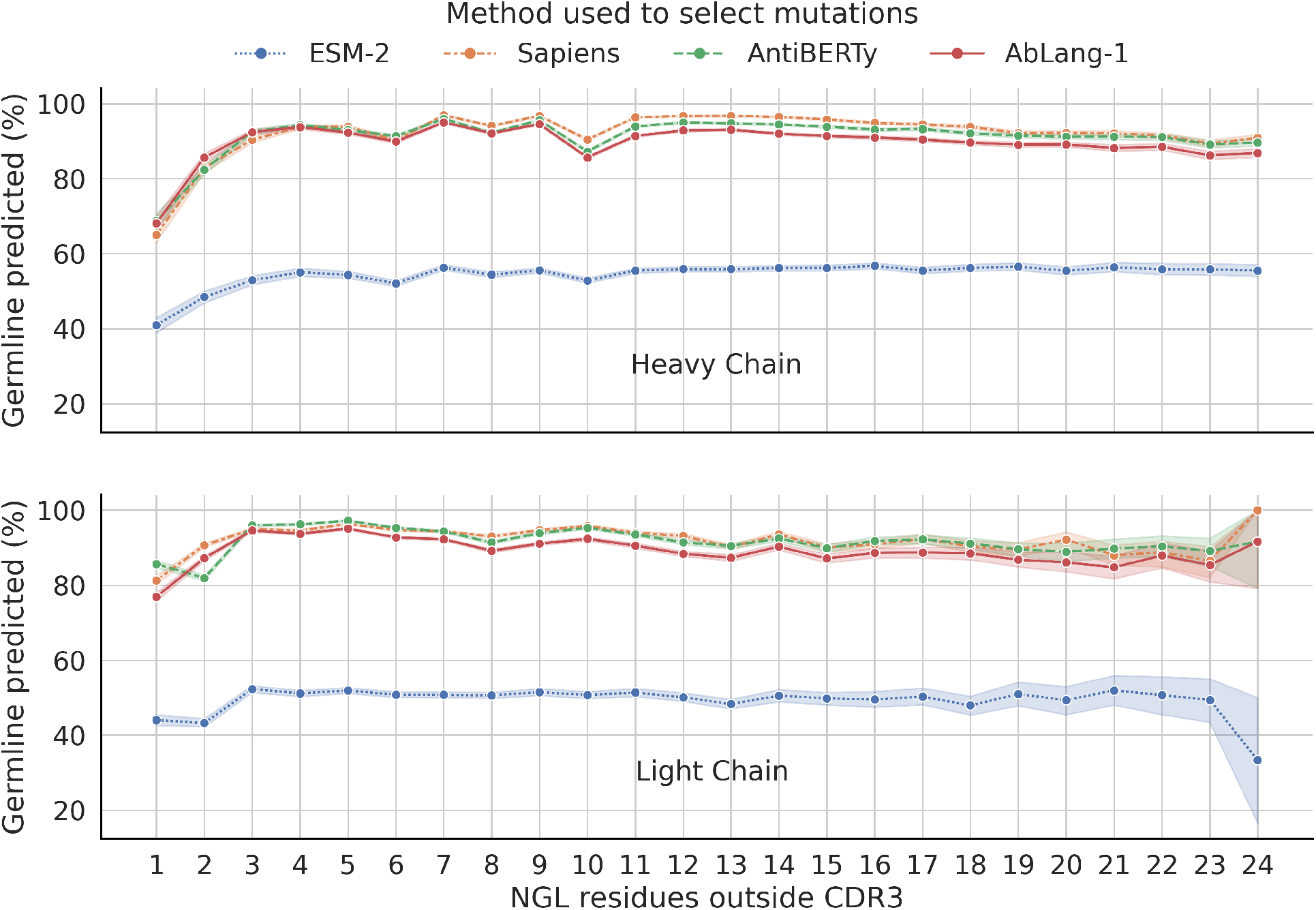
Germline prediction of masked non-germline (NGL) residues for four pre-trained language models (LMs). Results for the VH and VL are visualized separately, with predicted residues grouped by the number of NGL residues outside the CDR3 in their sequence. The 95% confidence interval is shown with the same colored error bands. The antibody-specific LMs Sapiens [21], AntiBERTy [22], and AbLang-1 [24] predict the germline 87.6%, 86.7%, and 84.9% of the time, respectively. ESM-2 [18], trained with few antibody sequences, predicts the germline 49.6% of the time. All LMs preferentially suggest mutations to the germline.

To investigate whether NGL residue predictions happen more frequently for sequences further from the germline, the results were grouped by the number of NGL residues per sequence (see Fig. 2). However, the frequency of germline predictions does not appear to be influenced by the number of NGL residues in a sequence. An exception is the slightly decreased germline prediction for sequences with only one NGL. This could be attributed to single nucleotide variants being wrongfully estimated as NGL.

To better understand what these models have learnt, we evaluated and compared their perplexity when predicting masked residues on three different sets. A set of random residues, representing the standard approach for calculating perplexity, a set of germline residues, and a set of NGL residues (see Methods 2.2). We calculated the perplexity for each set for ESM-2, AntiBERTy, and AbLang-1 (see Fig. 2). We left out Sapiens, as their predictions are similar to both AntiBERTy and AbLang-1. The perplexity metric spans from 1, denoting a perfect prediction, to positive infinity, representing zero probability for a correct prediction. The models have different vocabulary sizes ESM-2 (33), AntiBERTy (25), and AbLang-1 (24), but as they predict non canonical amino acids with close to zero probability, the best estimate of a random prediction is between the 20 canonical amino acids and would give a perplexity of 20.

Perplexity is normally calculated for all residues across the whole sequence or a random subset. With this standard approach, all models show a good performance at predicting masked residues (see Table 2). However, when evaluating specific regions, the performance on the more variable CDRs, especially CDR3, is considerably worse. As the CDRs only make up a small proportion of the residues in a chain the poor performance for this region is masked in the results for the whole chain.

**Table 2:** Perplexity comparison between the general-protein language model (LM) ESM-2 [18], and the antibody-specific LMs AntiBERTy [22] and AbLang-1 [24]. Perplexity was calculated on a set of randomly selected residues, only germline residues, and only non-germline (NGL) residues. Normally, perplexity is calculated for any residue across the whole sequence (red square), however; because of the relatively short length of the difficult to predict CDRs and the germline bias, the true performance for NGL residues is masked. The perplexity metric spans from 1, denoting a perfect prediction, to positive infinity, representing zero probability for a correct prediction. A random prediction would result in a perplexity of 20. Predictions worse than random are shown in *italic*.

|  | Residues | Heavy |  |  |  | Light |  |  |  |
| --- | --- | --- | --- | --- | --- | --- | --- | --- | --- |
|  |  | Whole | FRW | CDR1/2 | CDR3 | Whole | FRW | CDR1/2 | CDR3 |
| ESM-2 | All | 2.83 | 2.04 | 4.98 | 10.97 | 3.29 | 2.66 | 6.66 | 10.92 |
| AntiBERTy | All | 1.41 | 1.13 | 1.44 | 5.15 | 1.31 | 1.23 | 1.54 | 1.87 |
| AbLang-1 | All | 1.33 | 1.11 | 1.40 | 3.68 | 1.21 | 1.14 | 1.41 | 1.79 |
| ESM-2 | Germline | - | 1.91 | 4.12 | - | - | 2.54 | 6.11 | - |
| AntiBERTy | Germline | - | 1.05 | 1.10 | - | - | 1.17 | 1.28 | - |
| AbLang-1 | Germline | - | 1.03 | 1.08 | - | - | 1.07 | 1.16 | - |
| ESM-2 | Non-germline | - | *32.03* | *24.36* | *20.85* | - | *23.20* | *19.37* | *24.29* |
| AntiBERTy | Non-germline | - | *29.64* | *21.51* | *18.44* | - | *40.14* | *21.75* | *16.95* |
| AbLang-1 | Non-germline | - | *25.80* | *17.73* | *14.47* | - | *52.14* | *25.72* | *16.75* |

When further splitting residues into known germline and NGL residues, it becomes clear that the standard perplexity is heavily dominated by accurately predicting the germline and not the NGL residues. In fact, all models perform poorly at predicting NGL residues, while the antibody-specific LMs almost perfectly predict the germline residues. The germline bias also affects protein LMs, as seen with ESM-2’s poor NGL residue prediction. For all tested LMs, the prediction of NGL residues is close to or worse than random. Standard perplexity is not only skewed by the comparatively short length of CDRs, which hides the more difficult regions to predict, but also by the germline bias.

### 3.3 Reducing the germline bias

In an effort to reduce germline bias and improve NGL prediction, we trained several models while optimizing for NGL perplexity (see Methods 2.4). Starting from the architecture of a small ESM-2 model, each model introduced a new design choice. Table 3 shows each model and their incremental perplexity improvements. Our initial models Ab-Unpaired and Ab-Paired struggle with NGL prediction, like ESM-2, AntiBERTy, and AbLang-1. To focus training on NGL residues, cross-entropy loss was switched to focal loss, which heavily skews the loss towards poorly predicted residues. This significantly improves NGL predictions, going from perplexity values 14.23-38.95 to 10.24-12.69, without compromising germline accuracy.

Inspired by the idea of using a diverse set of pre-training objectives to train a model universally effective across downstream tasks [44], we modified the MLM approach to switch between random, short span, and long span masking, as well as dynamically changing the mask percentage. Using this modified masking technique slightly improved NGL perplexity in the framework and CDR1/2, but also mildly reduced performance in the CDR3.

Although pre-training on the large amounts of unpaired sequences (see Methods 2.1) before fine-tuning on paired sequences was expected to improve performance, as a more diverse set of sequences was seen during training, it led to a small dip in perplexity. This might be caused by the relative small number of training steps. As a final effort to optimize performance, we scaled the model from 6 to 12 layers and extended its training duration. This resulted in the best performing model, AbLang-2. All new models kept their near perfect prediction of masked germline residues, shown by their perplexity close to 1 (see Table 3).

**Table 3:** Perplexity comparison between the protein language model (LM) ESM-2 [18], the antibody-specific LMs AntiBERTy [22] and AbLang-1 [24], and our new selection of antibody-specific LMs (see Methods 2.4). While most of the models are near perfect at predicting masked germline residues, predictions for non-germline (NGL) residues show significantly higher perplexities. For ESM-2, AntiBERTy, AbLang-1, Ab-Unpaired, and Ab-Paired NGL perplexities are close to or worse than a random prediction. The largest improvement for NGL prediction came from switching to focal loss. Scaling up the model also improved performance, e.g. as seen by AbLang-2’s performances compared to Ab-FT.

|  | Germline residues |  |  |  | Non-germline residues |  |  |  |  |  |
| --- | --- | --- | --- | --- | --- | --- | --- | --- | --- | --- |
|  | Heavy |  | Light |  | Heavy |  |  | Light |  |  |
|  | FRW | CDR1/2 | FRW | CDR1/2 | FRW | CDR1/2 | CDR3 | FRW | CDR1/2 | CDR3 |
| ESM-2 | 1.91 | 4.12 | 2.54 | 6.11 | 32.03 | 24.36 | 20.85 | 23.20 | 19.37 | 24.29 |
| AntiBERTy | 1.05 | 1.10 | 1.17 | 1.28 | 29.64 | 21.51 | 18.44 | 40.14 | 21.75 | 16.95 |
| AbLang-1 | **1.03** | 1.08 | 1.07 | 1.16 | 25.80 | 17.73 | 14.47 | 52.14 | 25.72 | 16.75 |
| Ab-Unpaired | **1.02** | 1.07 | **1.01** | 1.05 | 26.81 | 18.95 | 14.42 | 37.60 | 19.37 | 17.25 |
| Ab-Paired | **1.02** | 1.06 | **1.02** | 1.05 | 27.24 | 18.70 | 14.23 | 38.95 | 19.25 | 16.98 |
| Ab-FL | 1.10 | 1.17 | 1.09 | 1.16 | 10.33 | 11.18 | 12.69 | 10.82 | 10.24 | 11.04 |
| Ab-ModMask | 1.11 | 1.18 | 1.09 | 1.17 | 10.26 | **11.13** | 13.18 | 10.78 | 10.19 | 11.42 |
| Ab-FT | 1.11 | 1.18 | 1.10 | 1.18 | 10.88 | 11.91 | 13.67 | 11.25 | 10.63 | 12.29 |
| AbLang-2 | 1.10 | 1.17 | 1.09 | 1.16 | **9.92** | **11.13** | **12.47** | **10.09** | **9.54** | **10.77** |

### 3.4 Clonotype mutations

The above perplexity is calculated with the presumption of a single correct prediction. In reality multiple amino acids are often a valid prediction at each position. To better verify the selection of suggested mutations, we examined positions in clonotypes with three or more known NGL residues. The clonotypes were created by grouping antibodies from the test set, based on identical source, V/J genes, and CDR3 length for both chains. This yielded 101 clonotypes, containing 226 and 60 sites with a minimum of three known NGL residues outside of the CDR3 in VHs and VLs, respectively. For each clonotype, a representative germline sequence was then generated by reverting NGL residues outside of the CDR3 back to the germline for the sequence with the fewest NGL residues.

For each site, the position was masked in the representative germline sequence and predicted using ESM-2, AntiBERTy, AbLang-1 and AbLang-2. The cumulative probability for known NGL residues at the site was then compared across the models (see Fig. 3a). For the VH, both AntiBERTy and AbLang-1 have an average cumulative probability below 2%, in contrast to ESM-2’s 20% and AbLang-2’s 15%. Similarly, for the VL, AntiBERTy and AbLang-1 have an average cumulative probability of 3% and 8%, respectively, while ESM-2 and AbLang-2 have 23% and 14%, respectively.

**Figure 3:**
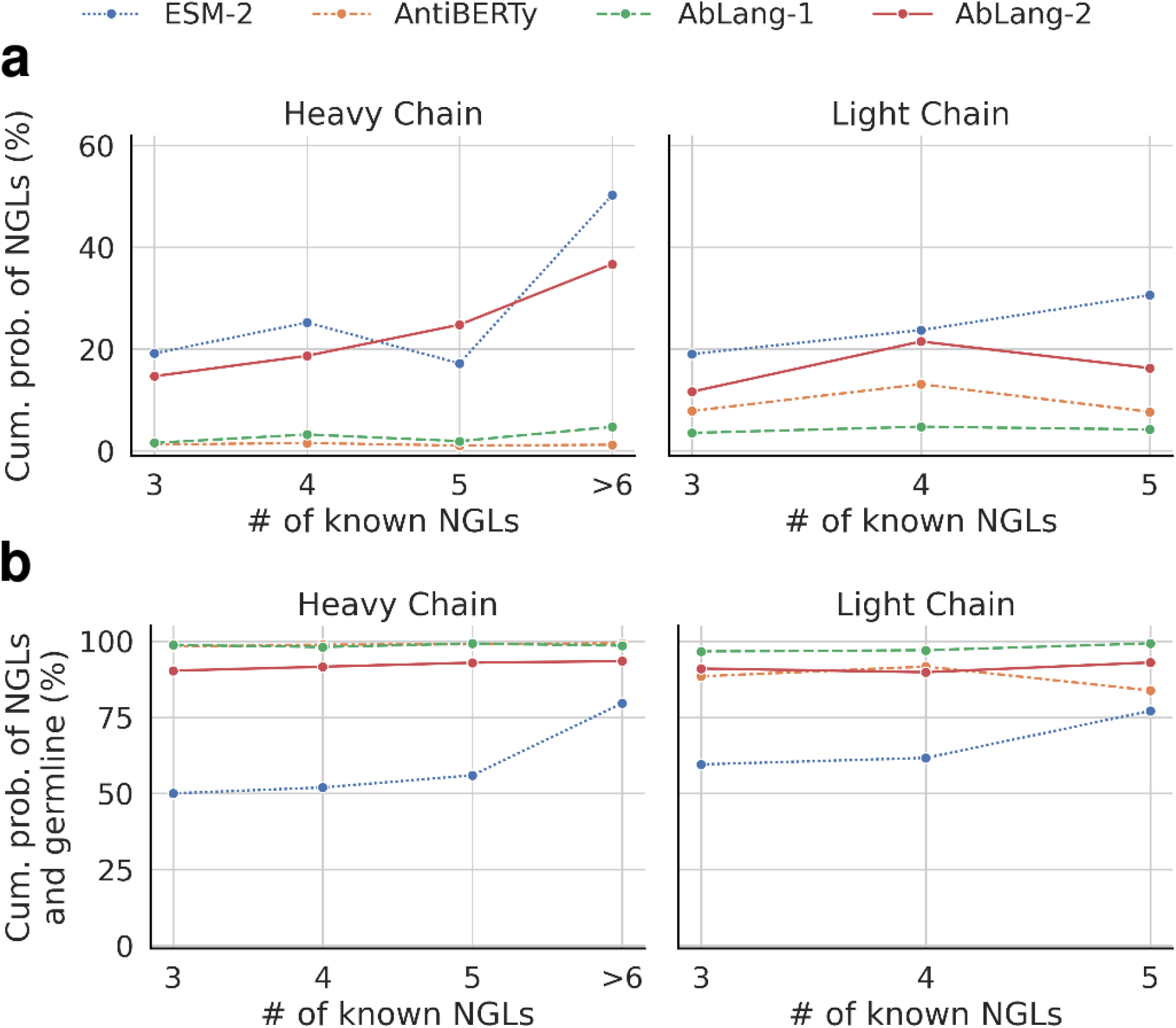
Comparison of cumulative probabilities of valid residues for the general protein language model (LM) ESM-2 [18] and the antibody-specific LMs AntiBERTy [22], AbLang-1 [24] and AbLang-2. Clonotypes were formed by grouping antibodies by source, V/J genes, and CDR3 length, yielding 101 clonotypes, containing 226 and 60 sites with a minimum of three known NGL residues outside of the CDR3 in VHs and VLs, respectively. **a**, Cumulative probabilities for known NGL residues. AntiBERTy and AbLang-1 show <2% for the VH, while ESM-2 and AbLang-2 display 20% and 15%. For the VL, values are 3% and 8% for AntiBERTy and AbLang-1, and 23% and 14% for ESM-2 and AbLang-2. **b**, Cumulative probabilities for known NGL residues and the germline. AntiBERTy, AbLang-1, and AbLang-2 demonstrate around 90-100% cumulative probabilities. ESM-2 presents 52% and 66% for the VHs and VLs, with the remaining probabilities suggesting amino acids different from the germline and at least three known NGL residues.

When the germline is included, see Fig. 3b, the cumulative probability for AntiBERTy, AbLang-1 and AbLang-2 hovers above 90%, underscoring how these models have a high probability of suggesting valid amino acids. In contrast, ESM-2 has a cumulative probability of 52% and 66% for the VH and VL, implying that ESM-2 potentially suggests invalid amino acids.

## 4 Discussion

Antibody sequences are predominantly composed of germline residues. Even those antibodies that are highly matured or have been optimized through extensive drug design campaigns, have only on average 15 and 20 NGL residues outside their CDR3s across both chains (see Fig. 1c). Over 93% of memory B-cells and 94% of therapeutics have five or more NGL residues across both chains. This suggests that while an extensive number of mutations away from the germline is rare, a select few are common for effective antibodies. While being able to suggest these mutations is vital for the design of therapeutic antibodies, identifying these specific mutations remains a significant challenge.

Pre-trained LMs like ESM-2, Sapiens, AntiBERTy, and AbLang are being used as a method to suggest potentially property enhancing mutations [25], so understanding the effects of the germline bias is important, as it limits the LMs ability to suggest relevant mutations. Our results demonstrate that all current LMs predominantly suggest germline residues and are poor predictors of NGL residues. The underperformance of these models in predicting NGL residues underscores the challenge of suggesting relevant changes to the germline.

To try and design an antibody-specific LM better capable of suggesting relevant NGL residues, we began with a small sized model with the same architecture as ESM-2 and iteratively improved it. First, the input was expanded to handle both unpaired and paired sequences. Then, focal loss was used during training, directing the model’s attention to the less represented NGL residues, resulting in improved performance of predicting NGL residues, as seen in Table 3. Drawing inspiration from the training of other LMs, we then modified the masking approach. To broaden the exposure to more diverse data, we first pre-trained the model on unpaired sequences before fine-tuning it on paired sequences. This allowed us to utilize the vast number of unpaired sequences, but still focus the model on handling paired data. For the final model, AbLang-2, we scaled up the size and training time resulting in our best model for predicting NGL residues.

Ideally, relevant mutations could be suggested directly from unmasked sequences, allowing a single forward pass instead of one for each masked residue. To measure the models’ capability to do this, we computed the perplexity when predicting NGL residues which had been reverted to the germline in the input (see Supplementary Table 1S). Although AbLang-2 shows the best performance, its perplexity is worse than random, highlighting the need for further work to enable LMs to suggest mutations away from an unmasked germline residue.

A problem with how we calculate perplexity is our presumption of a single correct prediction. However, for protein sequences, multiple mutations can be viable. While natural language has the same problem, natural LMs typically select from tens of thousands of unique tokens [46]. In contrast, protein-specific LMs choose from just 20 amino acids, with up to half sometimes being valid predictions. To better assess the models’ capacity to suggest valid mutations, we evaluated them on a dataset of same position mutations within clonotypes. AntiBERTy and AbLang-1 predict a known valid amino acid with >90% accuracy, however; they almost solely predict the germline (see Fig. 3). This limits their use for suggesting new relevant mutations. ESM-2 assigns higher probability to NGL residues, however; it also tends to predict mutations other than the known valid mutations (34%-48% of the time). AbLang-2 exhibits a high cumulative probability for NGL residues and simultaneously maintains a high probability for predicting known valid mutations. In other words, AbLang-2 suggests, with high probability, a diverse set of valid amino acids.

It is worth highlighting that we are only aware of a subset of the valid mutations and we do not weigh the potential importance of certain mutations over others. Moreover, as we predict masked residues from a representative germline, some NGL residues might not be viable within this sequence.

In this work, we demonstrate how the germline bias found in the OAS dataset which stems from the low ratio of non-germline mutations in both naturally occurring antibodies as well as highly optimized therapeutic antibodies, effects pre-trained LMs, especially how it affects their ability to suggest mutations away from the germline. In order to overcome this, we designed and pre-trained several antibody-specific LMs, with the final, AbLang-2, able to suggest a diverse set of valid mutations with high cumulative probability.

This work should facilitate the better design of therapeutic antibodies. For broader community engagement and research, we have made AbLang-2 freely and easily accessible via a python package (https://github.com/oxpig/AbLang2.git).

## Acknowledgements

This work was supported by the Engineering and Physical Sciences Research Council [EP/S024093/1]. This study received funding from GlaxoSmithKline plc.

For the purpose of Open Access, the author has applied a CC BY public copyright licence to any Author Accepted Manuscript (AAM) version arising from this submission.

## Supplementary Information

**Figure S1:**
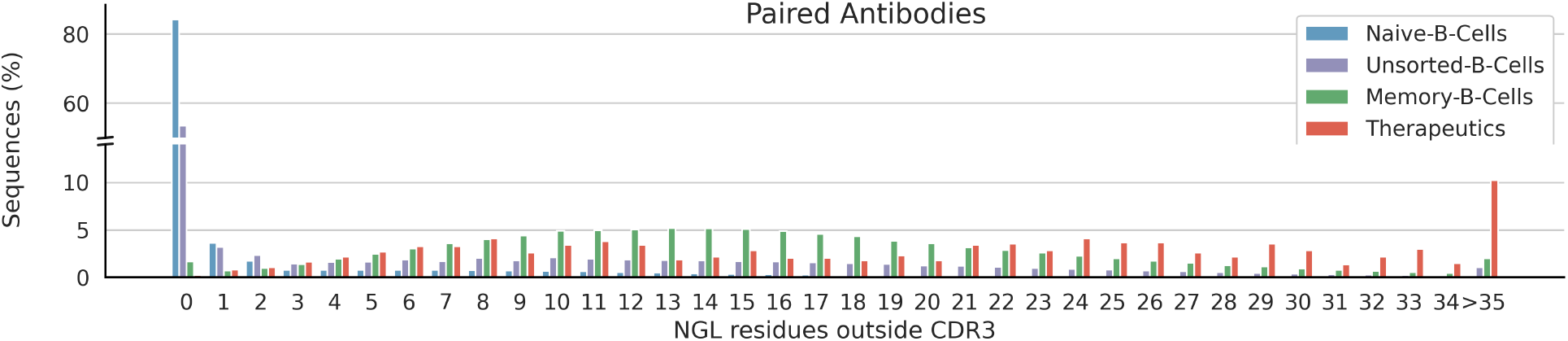
Distribution of NGL residues per VH-VL domain by source. Naive B-cell derived antibodies predominantly lack NGL residues, while memory B-cell derived antibodies display an average of *∼*15.3. Therapeutic antibodies exhibit an average of *∼*20.3 NGL residues.

**Table S1:**
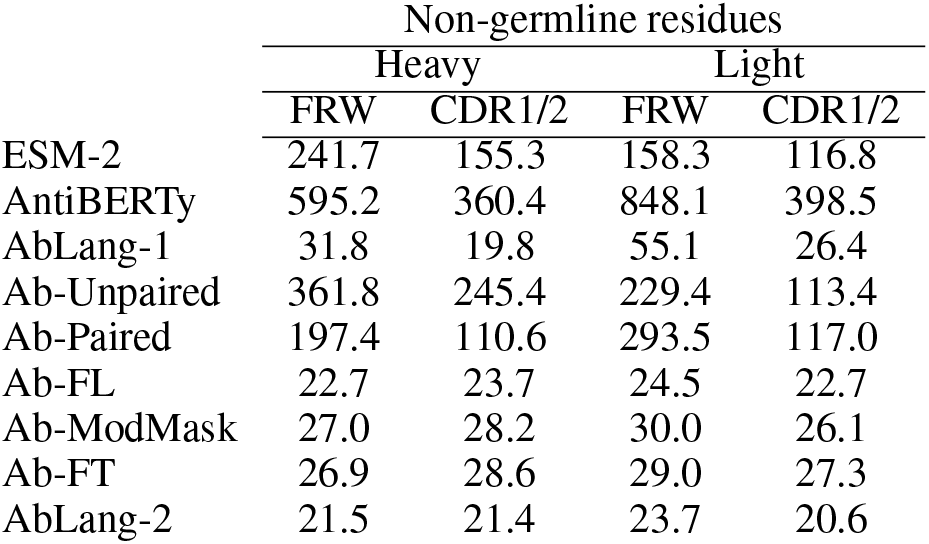
Comparison of perplexity computed when predicting NGL residues which has been reverted to the germline in the input. Comparison is between the general protein language model (LM) ESM-2 [18], the antibody-specific LMs AntiBERTy [22] and AbLang-1 [24], and our new selection of antibody-specific LMs (see Methods 2.4). Although AbLang-2 shows the best performance, its perplexity is only slightly better than random, highlighting the need for further work to enable LMs to suggest mutations away from a know germline.

|  | Non-germline residues |  |  |  |
| --- | --- | --- | --- | --- |
|  | Heavy |  | Light |  |
|  | FRW | CDR1/2 | FRW | CDR1/2 |
| ESM-2 | 241.7 | 155.3 | 158.3 | 116.8 |
| AntiBERTy | 595.2 | 360.4 | 848.1 | 398.5 |
| AbLang-1 | 31.8 | 19.8 | 55.1 | 26.4 |
| Ab-Unpaired | 361.8 | 245.4 | 229.4 | 113.4 |
| Ab-Paired | 197.4 | 110.6 | 293.5 | 117.0 |
| Ab-FL | 22.7 | 23.7 | 24.5 | 22.7 |
| Ab-ModMask | 27.0 | 28.2 | 30.0 | 26.1 |
| Ab-FT | 26.9 | 28.6 | 29.0 | 27.3 |
| AbLang-2 | 21.5 | 21.4 | 23.7 | 20.6 |

